# Size and shape regional differentiation during the development of the spine in the nine-banded armadillo (*Dasypus novemcinctus*)

**DOI:** 10.1101/2021.03.16.435620

**Authors:** Jillian D. Oliver, Katrina E. Jones, Stephanie E. Pierce, Lionel Hautier

## Abstract

Xenarthrans (armadillos, anteaters, sloths and their extinct relatives) are unique among mammals in displaying a distinctive specialization of the posterior trunk vertebrae - supernumerary vertebral xenarthrous articulations. This study seeks to understand how xenarthry develops through ontogeny and if it may be constrained to appear within pre-existing vertebral regions. Using 3D geometric morphometrics on the neural arches of vertebrae, we explore phenotypic, allometric, and disparity patterns of the different axial morphotypes during ontogeny of nine-banded armadillos. Shape-based regionalisation analyses showed that adult thoracolumbar column is divided into three regions according to the presence or absence of ribs and the presence or absence of xenarthrous articulations. A three-region-division was retrieved in almost all specimens through development, although younger stages (e.g. foetuses, neonates) have more region boundary variability. In size-based regionalisation analyses, thoracolumbar vertebrae are separated into two regions: a pre-diaphragmatic, pre-xenarthrous region, and a post-diaphragmatic xenarthrous region. We show that posterior thoracic vertebrae grow at a slower rate, while anterior thoracics and lumbar grow at a faster rate relatively, with rates decreasing anteroposterioly in the former and increasing anteroposterioly in the latter. We propose that different proportions between vertebrae and vertebral regions might result from differences in growth pattern and timing of ossification.

## INTRODUCTION

The vertebral column and its associated epaxial musculature function to maintain posture, propel the body, and act as an axis upon which limbs can move (e.g. Koob and Long, 2000). Across vertebrate lineages, the diversity of vertebral morphologies and patterning schemes reflects the breadth of modes of locomotion (e.g., Rockwell et al., 1938; Hebrank et al., 1990; Gal, 1993; Gál, 1993; Long et al., 1997; Boszczyk et al., 2001; Buchholtz, 2001; Schilling, 2011; Pierce et al., 2011; Werneburg et al., 2015; Molnar et al., 2015; Jones et al., 2021). Vertebral patterning, i.e. the regionalisation of vertebrae into distinct functional units, is implicated in the ecological and locomotor niches occupied by vertebrates (e.g.Rockwell et al., 1938; Buchholtz, 2007; Jones and German, 2014; Jones et al., 2018a, 2018b, 2020). In mammals, the prominence of asymmetric running gaits is facilitated by the division of the dorsal vertebrae into ribbed thoracic and rib-less lumbar regions. The thoracic region is further divided according to the diaphragmatic vertebra, which coincides with a shift in zygapophyseal morphology and thus intervertebral articulation.

The development of vertebrae proceeds from three ossification centres, the left and right vertebral arches and the centrum body, with two periods of vertebral growth being recognized during the development of the mouse vertebra (Bateman, 1954). The first phase corresponds to the fusion of the three ossification centres; the second phase consists of ’external accretion’ at the level of the vertebral arch, which grows outwards and upwards, coupled with ’internal erosion’ at the level of the neural canal (Bateman, 1954). On the centrum, endochondral ossification occurs at the epiphyses, and is mainly responsible for length increases. On the neural arches, periosteal ossification occurs at vertebral processes and is responsible for increases in height via appositional growth (Cubo et al., 2002; Valverde et al., 2010). By comparing vertebral development in different mouse strains, Johnson and O’Higgins (1994) suggested that early-developing features tend to show a more conserved growth pattern. They hypothesized that features showing the greatest specialization should complete their development later rather than earlier, with basic shared features of the vertebrae ossifying earlier than the more specialized features. Consequently, vertebral regionalization patterns might be more dynamic than previously conceived when looking at the full developmental sequence rather than adult morphology alone. However, no attempt has been made so far to characterize vertebral regionalization throughout ontogeny.

Xenarthrans (armadillos, anteaters, sloths and their extinct relatives) represent a special case among mammals for showing very specialized thoracolumbar series. In this group, the distinction between pre-diaphragmatic and post-diaphragmatic vertebrae is further defined by the presence of supernumerary vertebral articulations: the xenarthrous articulations or xenarthrales (Fig. 1). These articulations correspond to accessory intervertebral articulations formed between the anapophysis of a vertebra and the metapophysis of the vertebra immediately caudal to it, which span the posterior thoracic and lumbar regions and generally overlap with the post-diaphragmatic region (Gaudin, 1999; Oliver et al., 2016) (Fig. 1). Xenarthry is found in all modern and extinct xenarthrans with the exceptions of extant sloths and glyptodonts (Gaudin, 1999). It has been linked to evolution toward a fossorial lifestyle, the suggested ancestral locomotor mode of the lineage (Simpson, 1931; Frechkop, 1949; Jenkins, 1970; Gaudin and Biewener, 1992; Vizcaino and Milne, 2002; Nyakatura and Fischer, 2011; Emerling and Springer, 2015; Oliver et al., 2016; Olson et al., 2016). Recently, Oliver et al. (2016) demonstrated that xenarthrous vertebrae constitute a passively stiff region within the armadillo vertebral column, a functional attribute hypothesized to facilitate fossoriality by bracing the body, thereby providing leverage to the limbs during digging (Frechkop, 1949; Jenkins, 1970; Gaudin and Biewener, 1992). Despite this morphological uniqueness, few xenarthran studies have been interested in vertebral regionalisation or development (Gaudin and Biewener, 1992; MacPhee, 1994; Gaudin, 1999; Galliari et al., 2010; Nyakatura and Fischer, 2010; Galliari and Carlini, 2015), most of them focusing on the sloth departure from the cervical constant (Buchholtz and Stepien, 2009; Hautier et al., 2010; Varela-lasheras et al., 2011; Böhmer et al., 2018). In most cases, these studies only considered adult specimens and classically recognized mammalian vertebral regions (cervical, thoracic, lumbar, sacral, and caudal vertebrae).

**Figure 1.**
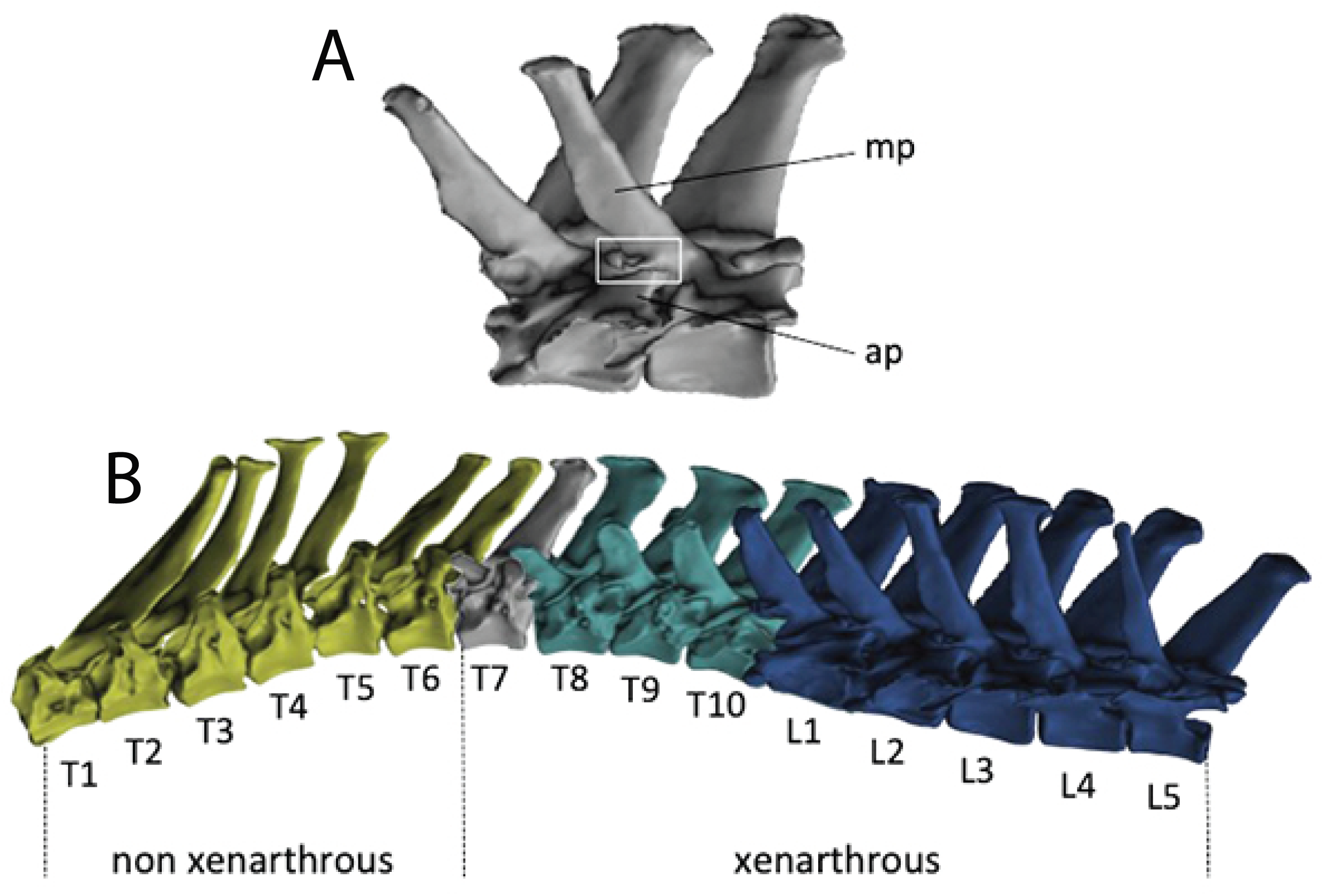
The xenarthrous articulation and motion segments of *Dasypus novemcinctus* used in experimentation. (A) The second and third lumbar vertebrae are shown in articulation, with the metapophysis (mp), anapophysis (ap) and xenarthrous articulation (xe) labeled. (B) The 10 thoracic vertebrae (T1–10) are subdivided into two regions: the anterior thoracic vertebrae (T1–6, yellow) are pre-xenarthrous and prediaphragmatic; the posterior thoracic vertebrae (T7–10, cyan) including the diaphragmatic vertebra (T7, grey) are xenarthrous and post-diaphragmatic. The lumbar vertebrae (L1-5, dark blue) are xenarthrous and post-diaphragmatic.

Developmental data has offered unique insights into the phenotypic trajectory of xenarthrous vertebrae, and invalidated the hypothesis that xenarthrous articulations represent a type of sacralization of the posterior thoracic and lumbar regions of xenarthran vertebral columns (Hautier et al., 2018). However, the development of the regionalisation patterns of their spine has never been thoroughly explored, while it could potentially explain how xenarthry has arisen within the context of the pre-existing vertebral regions. Here we ask if this apomorphic vertebral trait of xenarthrans may reflect a change in the regionalization patterns or if it may be constrained to appear within defined morphological, developmental and/or functional regions. We employ X-ray microtomography (X-ray µCT) and three-dimensional geometric morphometrics to describe vertebral shape at different developmental stages of the nine-banded armadillos, *Dasypus novemcinctus*. We use this data to explore phenotypic, allometric, and disparity patterns of vertebrae through development, and characterize vertebral regions of the xenarthran spine and how they develop. Based on preliminary works on regionalisation (e.g. Head and Polly, 2015; Randau and Goswami, 2017a; Jones et al., 2018b), we expect the thoracolumbar vertebrae of *D. novemcinctus* to be divided into three regions according to the thoracic/lumbar transition and the diaphragmatic vertebra. However, as vertebrae acquire diverging complex morphologies associated with specialized features through development (Johnson and O’Higgins, 1994), we also hypothesize that disparity and regionalisation should increase during ontogeny, concomitantly with the development of more specialized features such as xenarthrous articulations. Our unique morphometric dataset also enabled us to complement these regionalization and disparity analyses with comparisons of vertebral differential growth in order to extend our understanding of the ontogenetic development of the vertebral column.

## MATERIALS & METHODS

### Material

µCT scans of six *Dasypus spp*. foetuses, four neonates, two juveniles, and three adults were examined (Table 1). All specimens are stored in the collections of the Museum für Naturkunde Berlin (ZMB), the Smithsonian Museum of Natural History in Washington (USNM) and the Museum of Comparative Zoology in Harvard (MCZ), the Natural History Museum in London (BMNH), and the Association Kwata in Cayenne (France, JAGUARS collection, JAG). While individual absolute ages were unknown, specimens were chosen to cover an age range spanning the point at which all thoracolumbar vertebrae are ossified, through to adulthood, allowing us to examine the entire span of ossified vertebral development. All neonate specimens were labelled as such in the USNM records. The juvenile and adult specimens from the MZC were clearly identifiable as such prior to dissection. JAG-M1508 was directly compared to the known neonates and juveniles and fit it in the juvenile category accordingly. Relative ages of the foetuses were assessed based on degree of vertebral ossification (number and size of ossification centers on the entire vertebral column). We assumed that the extent of development of bony structures was a better indicator for age than size, due to substantial size intraspecific variation among *D. novemcinctus* (Hautier et al., 2017; Feijo et al., 2018). Average centroid sizes of left neural arches are, however, generally in line with this ranking, but some inconsistencies persist (Table 1). All individuals analysed were identified as *D. novemcinctus*, with the exception of ZMB 7b, whose species designation is unknown. However, we decided to include it in the dataset as its vertebral formula perfectly matches that of *D. novemcinctus*, but differs from other described *Dasypus* species (Galliari et al., 2010; Aya-Cuero et al., 2019). All specimens from the Smithsonian Museum of Natural History (USNM) and the Museum of Comparative Zoology (MCZ) were µCT-scanned on a SkyScan 1173 at the Museum of Comparative Zoology, Harvard University, and reconstructed with NRecon version 1.6.6.0 (MCZ scans were also included in Oliver et al., 2016). All other scans were taken prior to the commencement of this study (Hautier et al., 2010, 2011). Scans were segmented using either Avizo version 7.1 or Mimics Materialise software version 17.0.

**Table 1.** Specimens included in analyses. Specimens are listed in order of assumed increasing age, and are provided with average centroid sizes of the left neural arch, and their assigned stages. Specimens were aged according to extent of vertebral ossification, as observed in reconstructions of microCT scans. Average centroid sizes of left neural arches generally agree with this ranking, with inconsistencies due to intraspecific variation.

| Specimen ID | Museum ID | Average centroid size | Stage |
| --- | --- | --- | --- |
| A | ZMB 7b | 2.95 | Foetus |
| B | ZMB 40645 | 3.80 | Foetus |
| C | ZMB 40648 | 3.75 | Foetus |
| D | ZMB 40647 | 5.46 | Foetus |
| E | ZMB 85979 | 4.56 | Foetus |
| F | BMNH 2010_90_98_1 | 8.31 | Foetus |
| G | USNM 254678 | 6.92 | Neonate |
| H | USNM 204180 | 7.85 | Neonate |
| I | USNM 2960 | 7.59 | Neonate |
| J | USNM 21954 | 12.23 | Neonate |
| K | UM-M1508 | 11.05 | Juvenile |
| L | MCZ 67404 | 31.00 | Juvenile |
| M | MCZ 67407 | 40.88 | Adult |
| N | MCZ 67402 | 40.47 | Adult |
| O | MCZ 67401 | 38.47 | Adult |

### Geometric morphometrics

All specimens have ten thoracic vertebrae (T1-10) and five lumbar vertebrae (L1-5), with the exception of UM-M1508, which has ten thoracic and four lumbar vertebrae. In order to quantify the changes of shape across development, sixteen Type II landmarks were assigned to the left neural arch of all thoracolumbar vertebrae in each specimen with *MorphoDig* (Lebrun, 2014; Fig. 2). Analyses were restricted to neural arches as centra were absent in some cases or too poorly mineralized to enable performing morphometric comparisons. Landmarks were chosen to adequately capture the shape and the morphological transformations of the vertebrae (Table 2). Since most of the specimens analysed were developmentally immature, landmarks were assigned predominantly to structures identifiable in all vertebrae at all stages. Xenarthrous articulations ossify late during development (Hautier et al., 2018), so that no landmark could be placed directly onto them. Four landmarks (#8 to #11, Table 2) were placed on the metapophysis and the anapophysis. However, these two structures partially associated with xenarthry are for the most part present only in vertebrae participating in xenarthrous articulations, even if non-articulating anapophyses and metapophyses are often identified on T5 or T6 (Oliver et al., 2016). Since presence and absence are essential components of the morphological variation in the vertebral column, excluding them would lead to misleading results. To address the absence of these structures in a large subset of vertebrae sampled, we followed Klingenberg (2008), who suggested that in order to measure morphological novelty, landmarks should be assigned to the structure of interest when present, and to the position from which it would emerge when absent. This protocol has since been adapted for vertebral column, facilitating the quantification of morphologies present only in a subset of vertebrae under study (Head and Polly, 2015; Oliver et al., 2016; Jones et al., 2018b). We then performed geometric morphometric analyses on vertebral morphology to measure vertebral shape variance using the R package *geomorph* (Adams et al., 2020). All configurations (sets of landmarks) were superimposed using the Procrustes method of generalized least squares superimposition (GLS scaled, translated, and rotated configurations so that the intralandmark distances were minimized) and shape variability of the vertebrae was analysed by principal components analysis (PCA). A multivariate regression was performed to assess covariation patterns between the logarithm of the centroid size and Procrustes-aligned coordinates using procD.lm from *geomorph*. The hypothesis of parallel group slopes was assessed with a homogeneity of slopes (HOS) test. The HOS test includes pairwise comparisons between groups (vertebral regions) to assess significant differences of both the direction (angles) and magnitude (amount of change in shape with size) of allometric trajectories.

**Figure 2.**
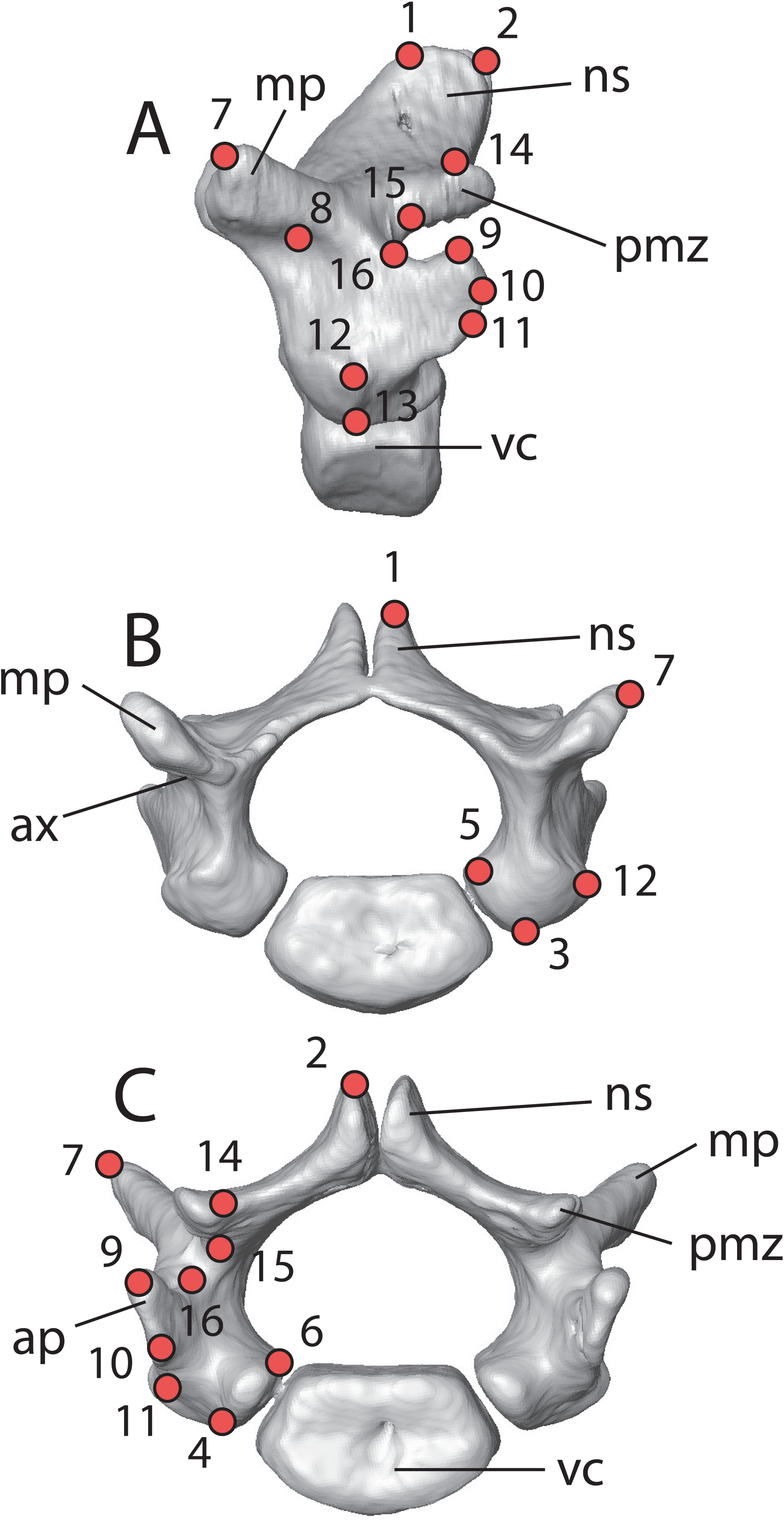
Landmarks digitized on the vertebrae. Lateral (A), anterior (B), and posterior (C) views. *Abbreviations*: ap, anapophysis; ax, anterior xenarthrous facet; mp, metapophysis; ns, neural spine; pmz, posterior medial zyg-apophysis; vc, vertebral centrum.

**Table 2.**
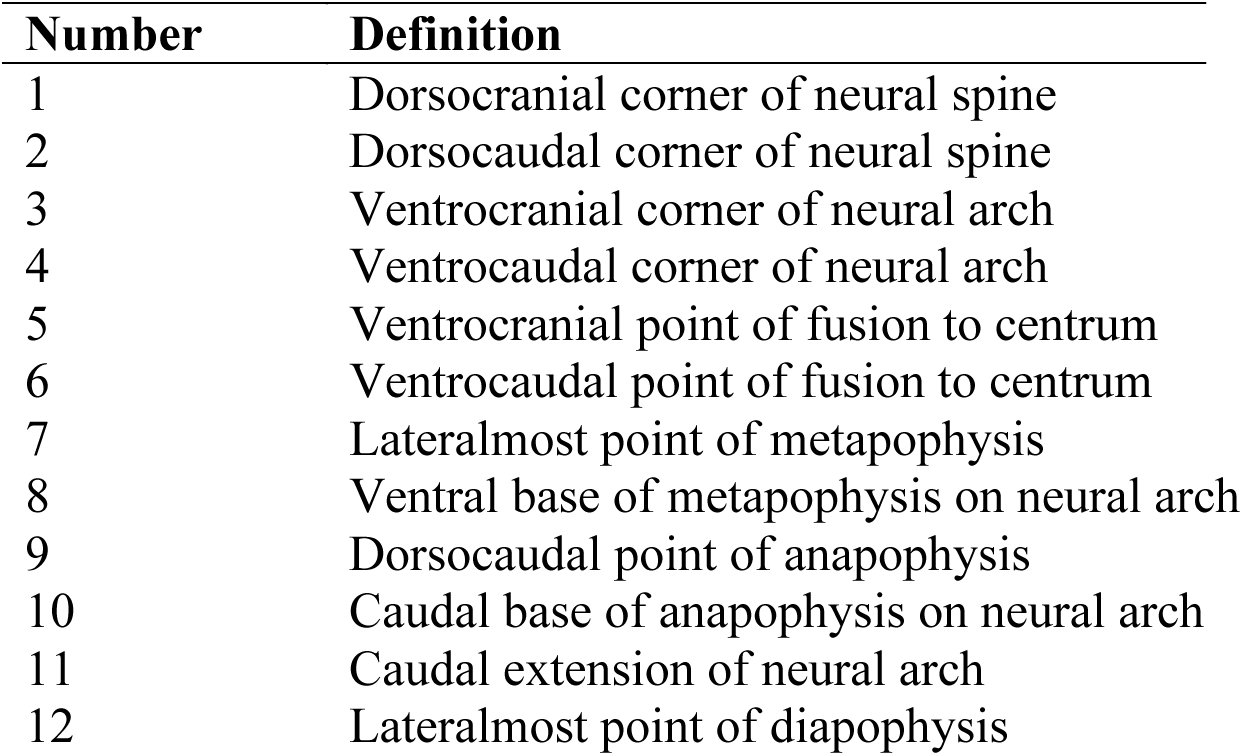
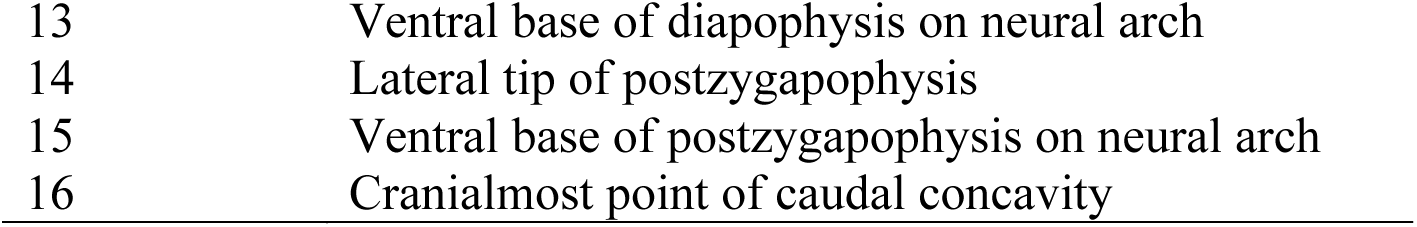
Locations of landmarks on the left neural arch.

### Regionalisation analysis

In order to examine the establishment of morphological regions during ossification of the vertebral column, we compared regionalisation patterns across developmental stages using the R package *Regions* (Jones et al., 2018a). We established region number and boundaries using a multivariate segmented regression approach that models regions as gradients, and maximum likelihood to select the optimal number of segments (Head and Polly 2015). For each specimen, every combination of regions was modelled iteratively, up to a maximum of six, based on either the Centroid Size (size analysis) or the top five PCs (shape analysis). The Akaike Information Criterion, corrected for small sample sizes, was used to determine the optimal regionalisation model (Jones et al., 2018a). The craniocaudal location of region breaks for each specimen was determined based on the position of the segmented regression boundaries in the best fit model. However, the curvilinear relationship between vertebral position and morphology introduces some error into boundary estimation, and thus boundary positions were considered accurate to within one vertebra (i.e., vertebral position +/- one).

### Phenotypic trajectory analyses

To determine the changes in regional gradient occurring through development, we used phenotypic trajectory analysis (PTA) (Adams and Collyer, 2009; Collyer and Adams, 2013). PTA enabled us to examine ontogenetic trajectories from two perspectives: across osteological development, as well as the morphological relationships between vertebrae within age groups. In both cases, phenotypic trajectories were drawn in a morphospace determined by the first two PC axes. In the across-column PTA, paths were drawn between region averages (i.e. anterior and xenarthrous thoracics, and lumbars) within each age group (foetus, neonate, juvenile, adult), indicating the amount of morphospace occupied by columns of different ages. In the PTA of ontogenetic trajectories, paths were drawn between regional averages of each age group, highlighting the diverging morphology of the recognized vertebral regions through development. Length, direction, and shape of trajectories were statistically compared pairwise in both analyses. The *geomorph* function trajectory.analysis (RRPP package) was used for both PTA’s (Adams et al., 2020).

### Integration and disparity analyses

To understand the relative evolvability of different regions we conducted integration and disparity analyses. Integration values of vertebrae across the column in every specimen were estimated as the standard deviation of the PCA eigenvalues accounting for 95% of shape variation (Young and Badyaev, 2006; Gómez-Robles and Polly, 2012). Greater dispersion of eigenvalues indicates stronger integration because it suggests more of the total variance is concentrated on just a few axes of variation. Following Gomez-Robles and Polly (2012), eigenvalues were standardized by total shape variance. This value provides a relative indication of the degree of morphological covariation between vertebrae in a single specimen, allowing us to assess the degree to which the vertebrae behave as a morphological unit. Conversely, values of disparity indicate the level of morphological diversity within the column, or within defined regions. These values allow us to assess the changes in morphological variability across developmental history. Disparity analyses were conducted using the *geomorph* function morphol.disparity, which uses Procrustes variation of a set of landmarks to measure pairwise morphological variance (Adams et al., 2020). Whole column disparity was calculated in each specimen, and regional (anterior thoracic, xenarthrous thoracic, lumbar) disparities were calculated in each age group (foetus, neonate, juvenile, adult). The assigned regions are in accordance with the adult regionalisation scheme predicted by the regionalisation analysis of centroid shape.

### Vertebral differential growth

To determine vertebral growth rates, we compared the centroid size of each vertebra to size of the thoracolumbar region, estimated by the sum of the centroid sizes of all of its constituting vertebrae (i.e anterior thoracics, xenarthrous thoracics, and lumbars). First, we estimated the proportion of each vertebra, as compared to size of the thoracolumbar region, during the four defined developmental stages (i.e foetuses, neonates, juveniles, and adults). We then performed multiple linear regressions of the centroid size of a vertebral locus against the size of the thoracolumbar region for all individuals of the ontogenetic series, and used the slope as an estimator of growth for each vertebra (Anderson and Secor, 2016).

## RESULTS

### Vertebral regionalisation

Regionalisation analyses were performed using centroid size and shape on thoracic and lumbar vertebrae from every specimen in order to determine the optimal number of regions in the vertebral columns across development. Shape-based regionalisation analysis revealed that thoracolumbar vertebrae were most often divided into three regions across all stages of development (Fig. 3A; Table 3). The vertebrae were divided into only two regions in two foetuses (specimens B and F), and into four regions in one neonate (specimen G). In the specimens in which the vertebrae were divided into three regions, the first break point ranged from T3-T6 (Table 3), which was situated at or anterior to the diaphragmatic vertebra, T7. The second breakpoint ranged from T7-T10 (Table 3), at or anterior to the thoracic-lumbar transition. In specimens B and F, the breakpoint between the two regions was T6 (Table 3). In specimen G, in which four regions were uncovered, the first two thoracic vertebrae formed their own region (Table 3). The three adults studied were found to have the same pattern of regionalisation, with breakpoints at T6 and T10 (Table 3), distinguishing a pre-diaphragmatic, pre-xenarthrous thoracic region (T1-6; hereafter referred to as anterior thoracics), a post-diaphragmatic xenarthrous thoracic region (T7-T10; hereafter referred to as posterior thoracics), and a lumbar region (L1-5). Adults consistently display higher values of Akaike weights than the juveniles, which indicate a better estimation of their best model.

**Figure 3.**
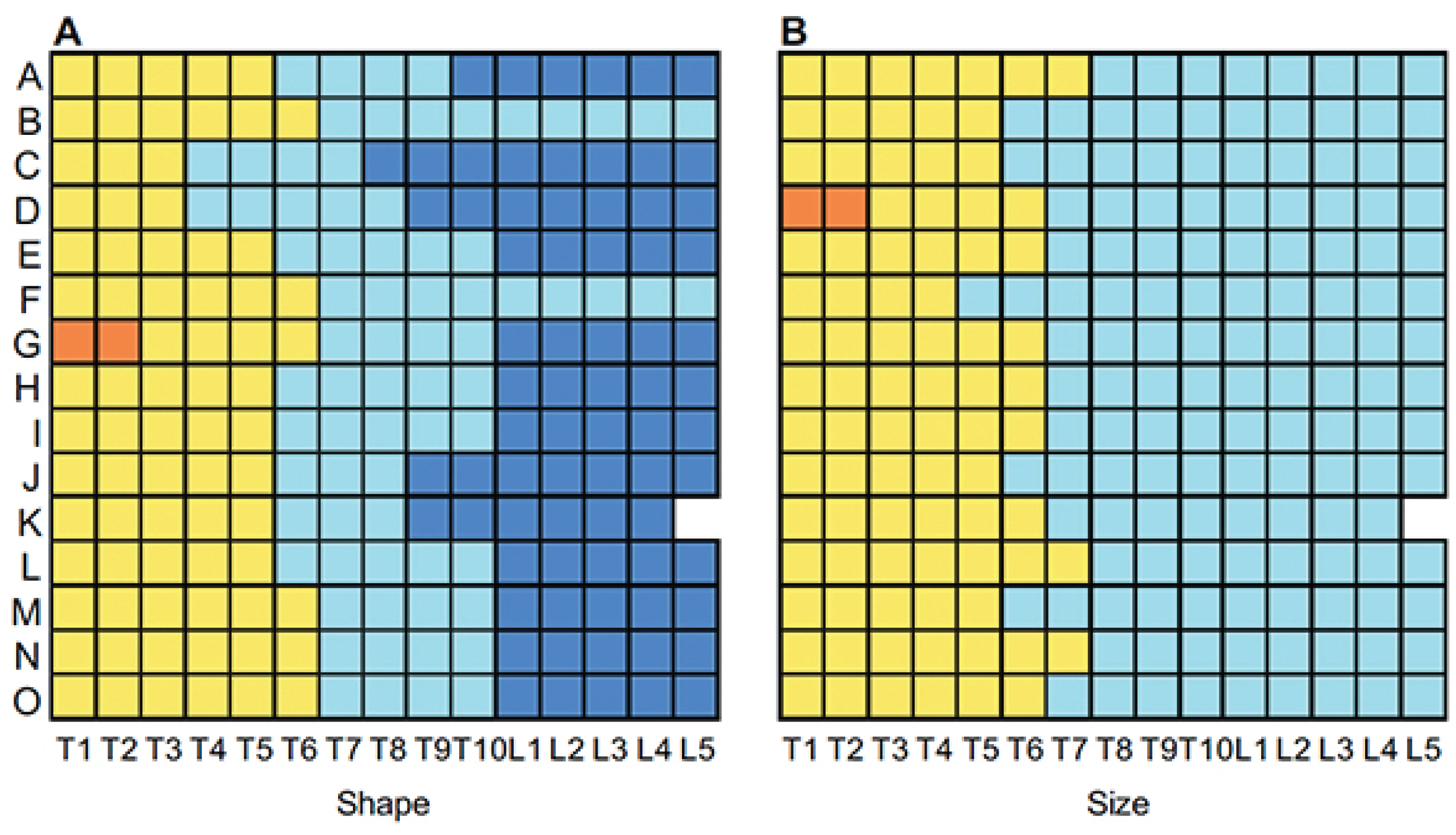
Vertebral regions predicted by (A) shape and (B) size, as identified according to the data presented in Table 2. Colors represent antero-posteriorly identified regions: yellow, anterior thoracics (i.e. pre-diaphragmatic non xenarthrous); cyan, posterior thoracics (i.e. post-diaphragmatic xenarthrous); dark blue, lumbars; orange, first two thoracics. Vertebral identities are denoted on the bottom of the figure, ranging from thoracic one (T1) to lumbar five (L5).

**Table 3.** Vertebral regions identified according to centroid shape and size. Regions identified according to shape are listed on the left, and regions identified according to size are on the right. Region score refers to the weighted mean derived from AIC analyses of regionalisation, which is rounded to identify the number of regions in the best model. Break points identify the locations within the thoracolumbar region separating the regions. Break point vertebrae are included in the anterior region, e.g. the three regions identified in Specimen A span from thoracic vertebrae (T) one to five, thoracics six to nine, and thoracic ten to lumbar (L) five.

| Specimen | Shape |  |  | Size |  |  |
| --- | --- | --- | --- | --- | --- | --- |
|  | Region score | Regions | Break point(s) | Region score | Regions | Break point(s) |
| A | 3.00 | 3 | T5, T9 | 2.12 | 2 | T7 |
| B | 2.15 | 2 | T6 | 2.00 | 2 | T5 |
| C | 2.86 | 3 | T3, T7 | 2.00 | 2 | T5 |
| D | 2.96 | 3 | T3, T8 | 2.91 | 3 | T2, T6 |
| E | 3.00 | 3 | T5, T10 | 2.03 | 2 | T6 |
| F | 2.20 | 2 | T6 | 2.00 | 2 | T4 |
| G | 3.83 | 4 | T2, T6, T10 | 2.21 | 2 | T6 |
| H | 2.94 | 3 | T5, T10 | 2.00 | 2 | T6 |
| I | 3.00 | 3 | T5, T10 | 2.01 | 2 | T6 |
| J | 2.98 | 3 | T5, T8 | 2.01 | 2 | T5 |
| K | 2.72 | 3 | T5, T8 | 2.09 | 2 | T6 |
| L | 3.00 | 3 | T5, T10 | 2.00 | 2 | T7 |
| M | 3.00 | 3 | T6, T10 | 2.05 | 2 | T5 |
| N | 3.00 | 3 | T6, T10 | 2.00 | 2 | T7 |
| O | 3.00 | 3 | T6, T10 | 2.00 | 2 | T6 |

Size-based regionalisation analysis settled on two regions for all specimens, with the exception of specimen D, in which the first two thoracic vertebrae formed their own region (Fig. 3B; Table 3). In specimens with two regions, the breakpoint ranged from T4-T7, with T6 being the most common (Table 3). Size-based regionalisation therefore roughly distinguished a pre-diaphragmatic, pre-xenarthrous region, and a post-diaphragmatic xenarthrous region, and does not differentiate between thoracic and lumbar vertebrae. In most cases, the location of the breakpoint was different between shape- and size-based regionalisation analyses, although it usually differs by only one vertebral position. No difference was observed in the values of Akaike weights between juveniles and adults.

### Phenotypic trajectories

The PCA performed on all individual vertebrae (Fig. 4A) confirmed that they present very similar morphology early in development. Developmental stages spread along PC1, which is mainly related to size, with foetuses and neonates occupying negative values and juveniles and adults the positive values. The first principal component is positively correlated with a dorsally projection of the neural spine (landmarks #1 and #2) and diapophysis (landmarks #12), an antero-medially projection of the metaphopysis (landmark #7), as well as more ventrally positioned postzygapophysis (landmarks #14 and #15) and anapophysis (landmarks #12 and #13) (Fig. 4A). In contrast, vertebral regions spread along PC2, which is positively correlated with a large antero-dorsal projection of the metaphopysis (landmark #7), a postero-lateral projection of the anapophysis (landmarks #12 and #13), and a lower neural spine (landmarks #1 and #2; Fig. 4A). Posterior thoracics (i.e. post-diaphragmatic xenarthrous) and lumbars are PC2 positive, and anterior thoracics are PC2 negative (Fig. 4A). Through ontogeny, we observed an increase in the amount of morphospace occupied by vertebrae, primarily along the second principal component (Fig. 4A). While vertebrae in foetuses are not readily distinguishable from each other, posterior thoracic and lumbar vertebrae take up gradually more morphospace through adulthood, allowing us to visualize the transition between vertebrae along the column along PC2 (Fig. 4A). In contrast, anterior thoracics occupy a distinct part of the morphospace and seem to individualize early during development, from neonates onwards, while they appear close to posterior thoracics in foetuses. In all cases, these three regions are well discriminated in all adults and one late juvenile, independently of their diaphragmatic or xenarthrous nature. The HOS test pairwise comparisons of ontogenetic allometric trajectories revealed no significant differences between the slopes of the different vertebral regions, but significant differences in magnitude. Vertebral morphospace defined by the third and fourth principal components (5.94% and 4.23% of the variance respectively), and associated morphological transformations, are presented in Fig. S1.

**Figure 4.**
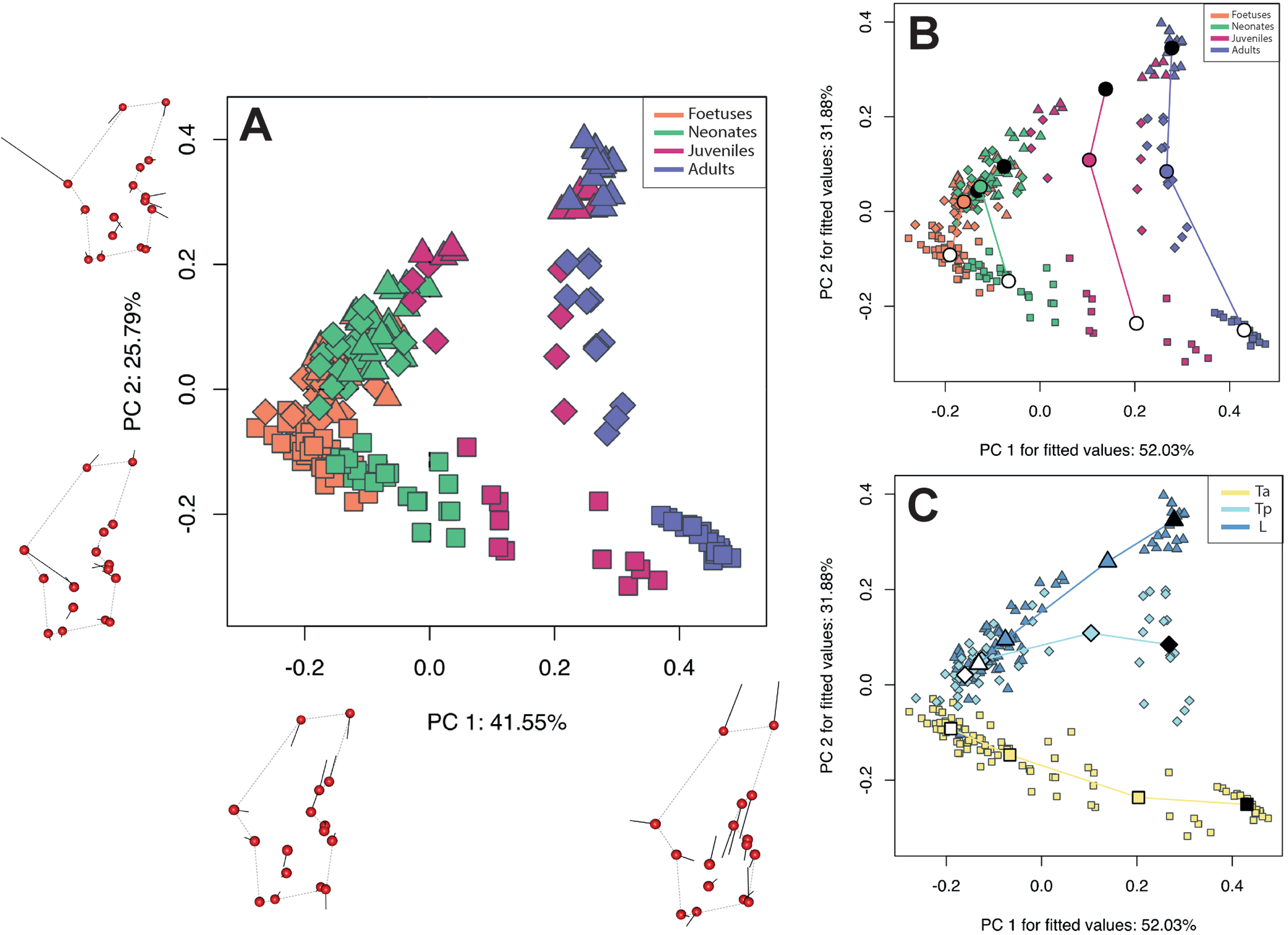
(A) Vertebral morphospace as identified by a principal component analysis and morphological variation expressed along the first two principal components. Dotted lines represent the outline of neural arches; vectors (black segments) underline the main directional changes. Symbol shapes characterize vertebral regions: squares are anterior thoracics, diamonds are xenarthrous thoracics, and triangles are lumbars. (B) Phenotypic trajectories are drawn between vertebral regions within each developmental stage. Symbol colors and shapes follow (A). Small coloured symbols denote individual vertebrae, and large coloured symbols denote the means of each vertebral region within each stage. The large white circles represent the means of anterior thoracic vertebrae in each stage, and the large black circles represent the means of lumbar vertebrae. (C) Phenotypic trajectories are drawn for each region across development. Symbol shapes follow (A). As in (B), small coloured symbols denote individual vertebrae, and large coloured symbols denote regional means within each developmental stage, with the exception of the large white symbols, which represent the mean of juvenile specimens, and the large black symbols which represent the mean of adult specimens.

Phenotypic trajectory analyses (PTAs; Fig. 4B and C) enabled us to examine in more detail this change in morphospace occupation across development. The first PTA traced trajectories between vertebral regions (anterior thoracics, posterior thoracics, and lumbars) in the four stages of development (Fig. 4B). The four developmental trajectories demonstrate a significant difference in size, as well as in direction except between juveniles and adults (Table 4). In contrast, they show no significant differences in shape (Table 4), the difference being marginally significant between foetuses and adults. The second PTA defined trajectories within regions across developmental stages and demonstrates the diverging morphological paths taken by the three regions through development (Fig. 4C). Posterior thoracic and lumbar vertebrae display very similar trajectories at the beginning of the ontogenetic sequence, then gradually spread out to occupy distinct morphospaces in adulthood (PC1 positive). The three regional trajectories are significantly different from each other in direction (Table 5), but show no significant difference in shape. Anterior thoracics show significant differences in size trajectories with both posterior thoracics and lumbars, which in turn do not significantly differ from each other (Table 5).

**Table 4.** Pairwise comparisons between phenotypic trajectories of the vertebral regions between developmental stages. Significant differences and *p*-values according to *p*≤0.05 are bolded.

| | Size<br>$P$ value | Direction<br>$P$ value | Shape<br>$P$ value |
| --- | --- | --- | --- |
| Foetus x Neonate | **0.004** | **0.001** | 0.873 |
| Foetus x Juvenile | **0.001** | **0.001** | 0.565 |
| Foetus x Adult | **0.001** | **0.001** | 0.081 |
| Neonate x Juvenile | **0.001** | **0.001** | 0.625 |
| Neonate x Adult | **0.001** | **0.001** | 0.157 |
| Juvenile x Adult | **0.041** | 0.118 | 0.518 |

**Table 5.** Pairwise comparisons between phenotypic trajectories of the developmental stages between vertebral regions. Significant differences and *p*-values according to *p*≤0.05 are bolded.

| | Size<br>$P$ value | Direction<br>$P$ value | Shape<br>$P$ value |
| --- | --- | --- | --- |
| Ta x Tx | **0.001** | **0.001** | 0.054 |
| Ta x L | **0.002** | **0.001** | 0.181 |
| Tx x L | 0.143 | **0.001** | 0.847 |

### Vertebral disparity and integration

The whole-column disparity and integration were calculated for every specimen (Fig. 5A), and regional disparity was then calculated across developmental stages (Fig. 5B). Whole-column integration gradually increased through prenatal development, and rapidly increased in post-natal development (Fig. 5A). Whole-column disparity initially decreased from the foetus stage to the neonate stage, and then increased again in juveniles and adults (Fig. 5A), indicating that early development is characterized by a stage of relatively high morphological variability in vertebrae, while morphologies appear to converge in the neonatal stage. As with integration, disparity rapidly increased postnatally to reach a peak in adulthood.

**Figure 5.**
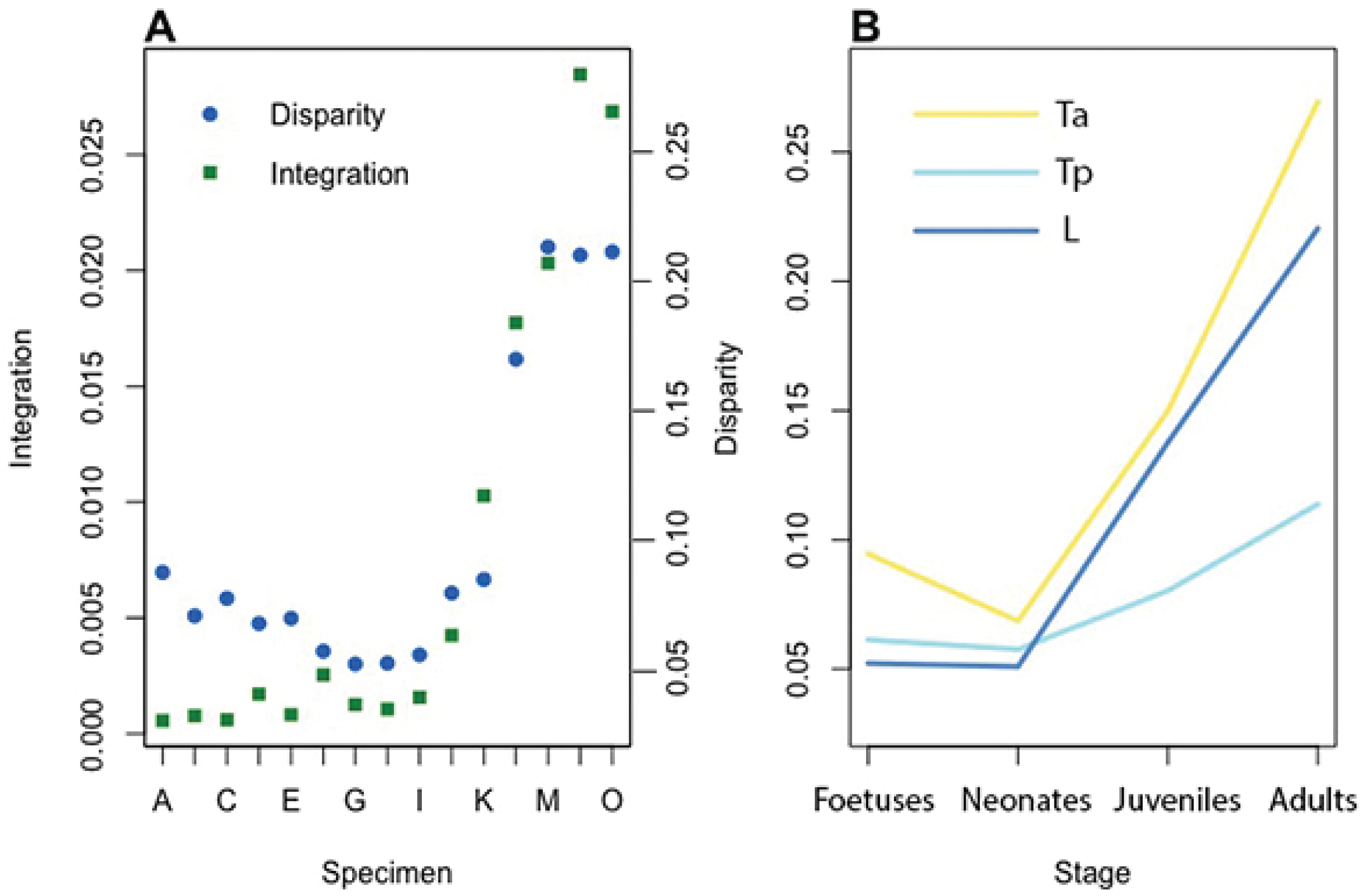
Morphological disparity and integration across development. In (A), whole-column disparity and integration are plotted across individuals, in order from youngest to oldest. In (B), regional disparity is plotted across the four developmental stages.

Examining with-in regional disparity across stages of development allowed us to further dissect the disparity data (Fig. 5B). In all three regions, as defined by regionalisation analyses, disparity significantly increased across development. The high disparity observed in adult vertebral columns was primarily localized to anterior thoracic (Ta) and lumbar (L) regions, which were significantly more disparate than they were in the three earlier stages of development (Table 6). A more modest increase in disparity is observed in the posterior thoracic (Tp) region in adulthood, which was significantly different only from foetus and neonate disparities (Table 6). Foetus and neonate disparities were not significantly different from each other in any of the three regions (Table 6), despite a clear drop in disparity from the foetus to neonate stage in the anterior thoracic region (Fig. 5B). This decrease mimics that seen in whole-column disparity values, indicating that this pre- and neonatal drop in disparity is likely localized to the anterior thoracic region.

**Table 6.** Pairwise differences of regional morphological disparities across developmental stages. Absolute pairwise difference between disparities are presented in the top right of each region, and *p*-values of the corresponding comparisons are presented in the bottom left. Significant differences and *p*-values according to *p*≤0.05 are bolded.

|  |  | Foetus | Neonate | Juvenile | Adult |
| --- | --- | --- | --- | --- | --- |
| Ta |  |  |  |  |  |
| Ta | Foetus | / | 0.0260 | **0.0554** | **0.1749** |
|  | Neonate | 0.1717 | / | **0.0814** | **0.2009** |
|  | Juvenile | **0.0206** | **0.0012** | / | **0.1196** |
|  | Adult | **0.0001** | **0.0001** | **0.0001** | / |
| Tx |  |  |  |  |  |
| Tx | Foetus | / | 0.0037 | 0.019 | **0.0525** |
|  | Neonate | 0.8699 | / | 0.0227 | **0.0562** |
|  | Juvenile | 0.5261 | 0.4946 | / | 0.0335 |
|  | Adult | **0.0398** | **0.0388** | 0.307 | / |
| L |  |  |  |  |  |
| L | Foetus | / | 0.0013 | **0.0855** | **0.1683** |
|  | Neonate | 0.952 | / | **0.0868** | **0.1696** |
|  | Juvenile | **0.0018** | **0.0021** | / | **0.0823** |
|  | Adult | **0.0001** | **0.0001** | **0.0053** | / |

### Regional axial growth

Vertebral proportions are very similar between ontogenetic stages (Fig. 6A), but two trends are noticeable: the mid-trunk region becomes relatively shorter, while most growth seems to occur toward the anterior and posterior extremes of the thoracolumbar region. The slope of the linear regressions of the centroid size of each vertebra against the size of the whole thoracolumbar region enabled us to estimate the growth patterns for each vertebra (Fig. 6B). Growth rates steadily decrease from the first thoracic vertebra to the first xenarthrous thoracic vertebra, to increase again up to the fourth lumbar vertebra, the last lumbar showing a slower rate than the vertebra just anterior to it. In sum, posterior thoracic vertebrae seem to grow at a slower rate, while anterior thoracics and lumbar grow at a faster rate relatively (Fig. 6B). The rate decreases anteroposterioly in anterior thoracics, but increases anteroposterioly in lumbars.

**Figure 6.**
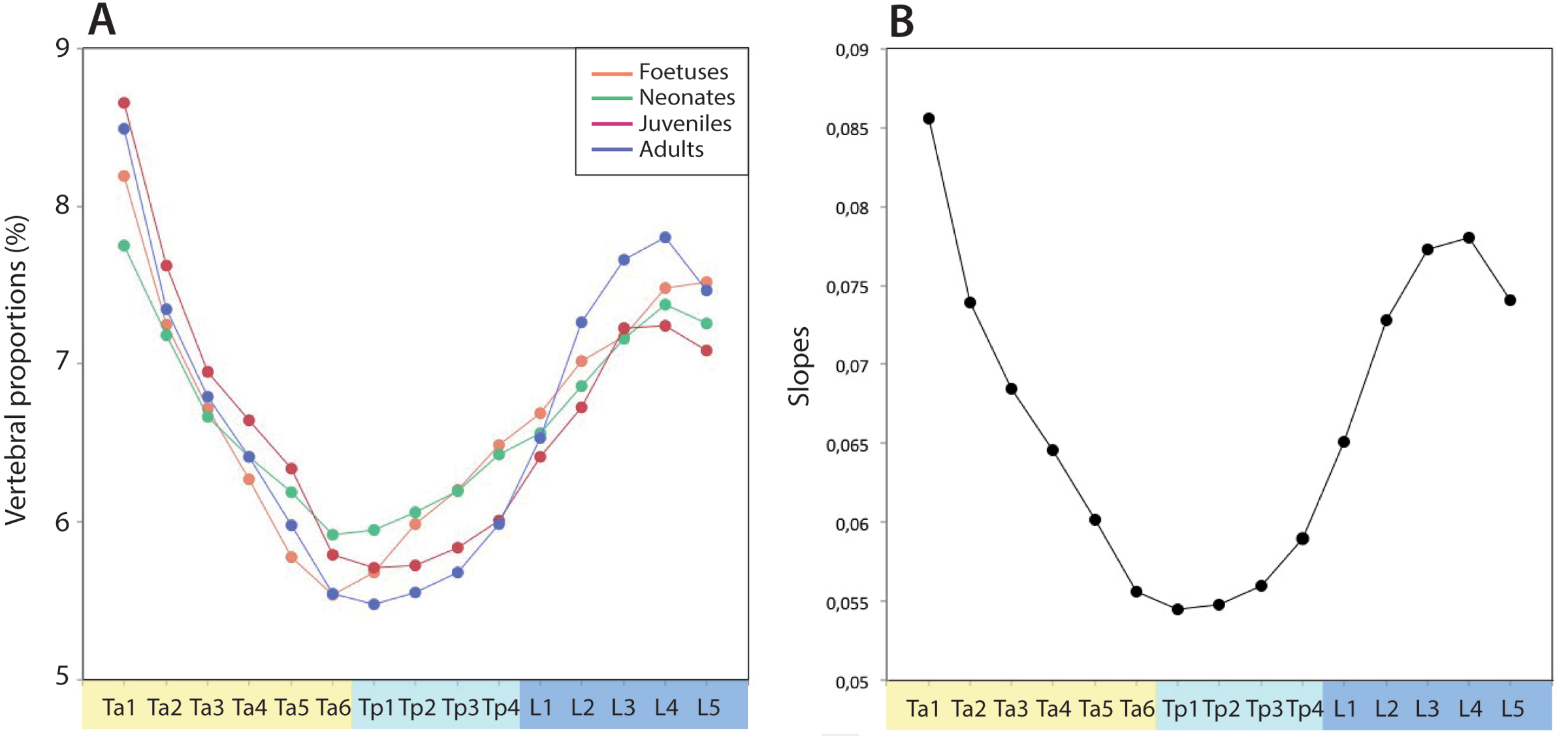
(A) Proportion of each vertebra, as compared to size of the thoracolumbar region during the four developmental stages (i.e foetuses, neonates, juveniles, and adults) (B) Regression slopes plotted for the centroid of each vertebral position against the size of the thoracolumbar region for all individuals. The slope is then used as an estimator of vertebral growth.

## DISCUSSION

### Regionalisation developmental dynamics

Qualitative examination of the *Dasypus* thoracolumbar region reveals that it is a morphologically complex series with gradual intervertebral anatomical transition, and that foetuses and juveniles display more homogeneous vertebral columns than the adults with little observable regional differentiation (Hautier et al., 2018). These qualitative observations were confirmed here by our quantitative analyses, with the thoracolumbar vertebral series becoming more consistently regionalised as *Dasypus* develops into its adult form. Although no landmark could be placed directly onto all zygapophyseal facets and xenarthrous articulations, the diaphragmatic vertebra clearly marks the transition between anterior and posterior thoracics. This implies that this modular pattern involves the whole neural arch morphology, and that it precedes the establishment of the shift in zygapophyseal morphology and intervertebral articulations. In most foetuses and pre-adults, the semi-xenarthrous vertebra (T6) is grouped with posterior thoracic vertebrae, suggesting that its shape is predominantly defined by its posterior xenarthrous articulation. This may also be a product of either the reduction of the metapophysis (landmark #7) in a posterior-to-anterior direction (from L5 to T7) in the adult (Gaudin, 1999), or of later ossification of the mid-thoracic vertebrae as compared to the anterior thoracics and lumbars (Hautier et al., 2010 and Fig. 7). As the development of the vertebral column proceeds, T7-T10 consistently group in posterior thoracics at a moment when the xenarthrous characteristics of T7, and of the succeeding vertebrae, become more pronounced (Hautier et al., 2018).

**Figure 7.**
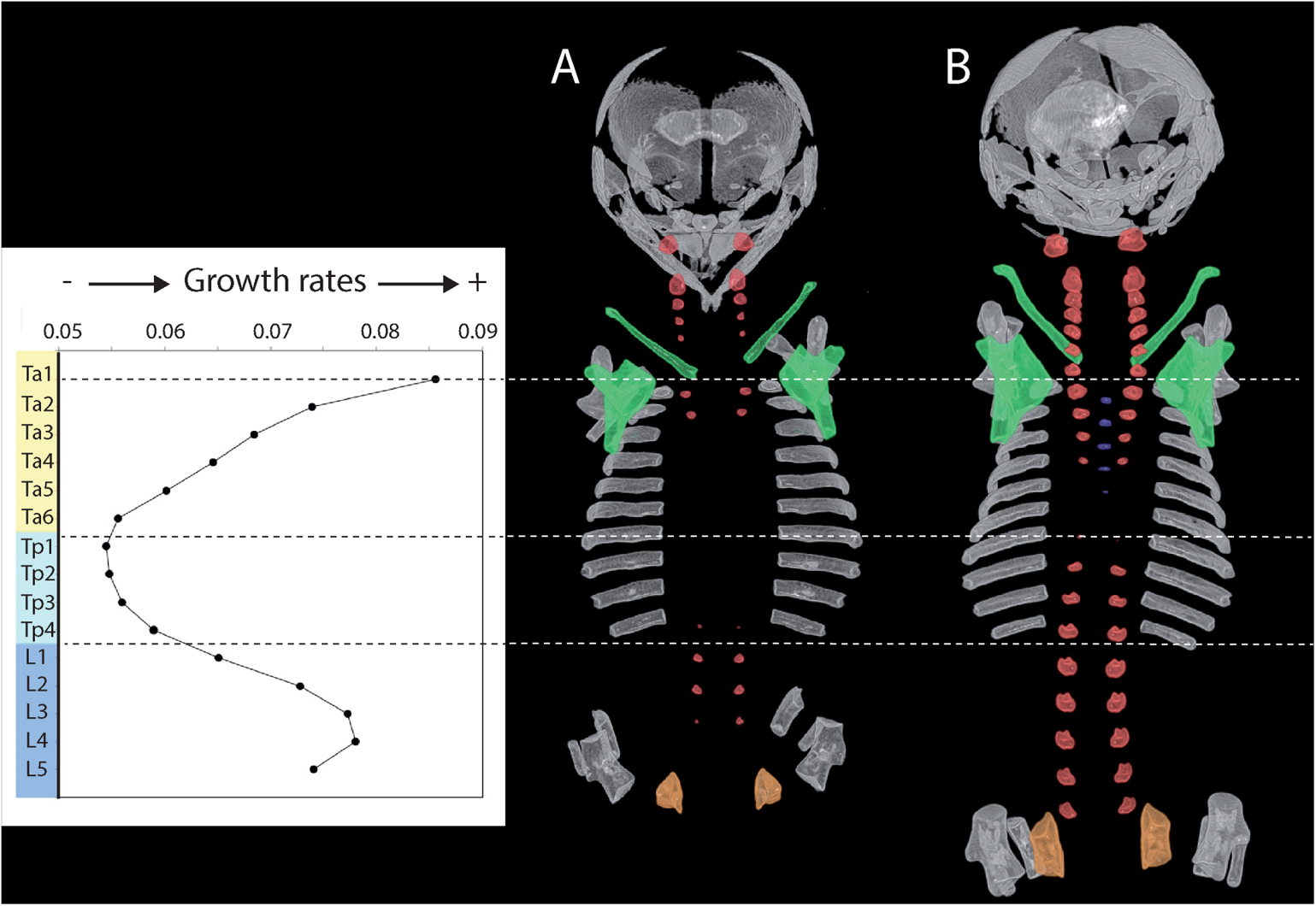
Ossification sequences in the long-nosed armadillo (*Dasypus*) vertebral elements as compared to vertebral growth. Dorsal view of the rib cage in two armadillo foetuses (A and B), following Hautier et al. (2010). Vertebral neural arches are in red, vertebral centra are in blue, scapula and clavicle are in green, and ischium, ilium, and pubis are in orange. For clarity, elements of the zeugopod have been removed.

Unlike the consistent boundaries of the adult specimens, regional transitions in foetuses and stillborn appeared more variable (Fig. 2). This variability in boundary definitions could be attributable to imprecisions in age determination or to more variable degrees of ossification in young specimens, influencing landmark placement. Such inconsistency might also result from imprecisions in boundary placement resulting from more subtle or more gradational transitions between regions. The approach applied here models the vertebral column as a segmented regression composed of a series of linear relationships (Head and Polly 2015). This model provides a better approximation in the adult specimens where the regions are clearly defined. Whereas, more curvilinear patterns in younger specimens make determining regional patterns more challenging. However, in shape-based regionalisation analyses, a three-region-division was obtained in almost all specimens (i.e. 12 out of the 15 specimens). The main exceptions are two foetuses in which only two regions were recognized, with a break point located at the diaphragmatic vertebra, and one foetus with four regions, in which the first two anterior thoracic vertebrae form their own region. This result suggests that axial shape regionalisation is conserved during development despite drastic changes in vertebral morphology. The distinction between anterior and posterior thoracic vertebrae rapidly becomes more discrete, as exemplified in the two foetuses for which the thoracolumbar vertebrae are divided into two groups (Fig. 4B). Such results indicate that differentiation of vertebral regions in the vertebral column of xenarthrans is not directly coincident with the ontogenetic development of the specialized xenarthrous joints, which only start to ossify at the neonate stage (Hautier et al., 2018), but instead occurs very early on during development when the shape of vertebrae remains quite simple. Most importantly, this two-region model likely reflects a subtler distinction between thoracics and lumbars than between pre-diaphragmatic and post-diaphragmatic vertebrae. This was confirmed by the PTA results, with posterior thoracic and lumbar vertebrae displaying more similar trajectories (Fig. 4C).

As is often the case in morphological evolution, allometry is likely to explain a substantial portion of the morphological differentiation of the spine, as size increases need to be adjusted by structural adaptation to static loading (Randau et al., 2016; Jones et al., 2018b). Our developmental data confirmed this in showing the significant influence of size on the regionalisation of the axial skeleton. In size-based regionalisation, two main regions can be recognized through development (Fig. 3B), with the break point located around the diaphragmatic vertebra. In one young foetus, however, the first two thoracic vertebrae form their own group again. This might be attributed to the fact that these two neural arches ossify earlier than the more caudal vertebrae (Hautier et al., 2010; Fig. 7), and are thus larger to a variable extent. Apart from a few specimens, the diaphragmatic vertebra was consistently attributed to the posterior thoracic region whereas T6 inconsistently fits either of these two primary regions, probably as it displays transitional morphological characters reflective of both anterior and posterior thoracic vertebrae, including semi-xenarthrous articulations. Although intuitive, this size-based pattern cannot be attributed entirely to the presence of large metapophyses in posterior thoracic and lumbar vertebrae, because such protrusions are not developed until late during development (Fig. 4A). In the youngest foetuses, the metapophyses can only be identified as a small rounded bump, but become more and more distinguishable through development on the lumbar series, the posterior thoracics following the same trajectory later during development (Hautier et al., 2018). Interestingly, the retrieved two-region model based on size, and on shape to a lesser extent (specimen B and F, Fig. 3A), does not take into account the anatomically and functionally relevant distinction between the rib-bearing thoracic vertebrae and rib-less lumbar vertebrae. Such division might instead be linked to the development of the sternum, since post-diaphragmatic thoracics are the origin for the floating ribs in mammals, which are not connected to the former. Consequently, the distinction between anterior and posterior thoracic regions might relate more to ribs than vertebrae themselves, and be reflective of their different function.

The mammalian vertebral column is commonly organized into cervical, thoracic, lumbar, sacral, and caudal regions according to a common *Hox* expression scheme, and to the functional relevance of each region (Burke et al., 1995; Wellik, 2007). The thoracolumbar regionalisation scheme described here in *Dasypus* reaches beyond what is typically suggested for mammals, notably for xenarthrans in which only classical vertebral regions were previously recognized and discussed (Gaudin, 1999; Buchholtz and Stepien, 2009; Galliari et al., 2010; Hautier et al., 2010; Nyakatura and Fischer, 2010; Varela-lasheras et al., 2011; Galliari and Carlini, 2015). The thoracic region is here divided into two morphological and possibly functional segments, the most posterior of which is identified both by articulation with ribs, and by the presence of post-diaphragmatic zygapophyseal and xenarthrous articulations. Our size-based regionalisation analyses showed that this distinction between anterior and posterior thoracics might be largely influenced by allometry, so that growth pattern may constitute a key feature underlying regionalisation differences. In terms of morphology alone, subdivision of thoracic vertebrae into two regions was recently proposed for *M. musculus* and a variety of other amniotes (Head and Polly, 2015; Jones et al., 2018a), although the placement of the subdivision was taxon-specific and appears to correlate with the *Hox* code. Rather than a novel trait unique to xenarthrans, the additional regionalisation found in *Dasypus* may then constitute a shared feature of all mammals, and perhaps amniotes, and could be considered as an exaptation of an existing region to pattern a new type of morphology. This proposition concurs with the conclusions of Woltering and Duboule (2015) who proposed that morphological novelties in the axial skeleton could coincide with pre-existing modules and represent an exaptation of the axial *Hox* pattern.

### Ontogenetic disparity and integration

Previous studies (Bergmann et al., 2006; Jones and German, 2014) proposed that functional demands on the vertebral column differ prenatally, neonatally, and in adulthood. Naturally, neonates require integrated vertebral columns as much as juveniles and adults, but ranges of axial movements might be more limited at younger stages. The whole-column integration was then expected to increase across development of ossified parts, especially postnatally, as vertebral function and mobility are dependent on the individual units working in tandem. This trend was observed across our specimens, but with a dramatic increase in juveniles and adults. Although precocial, nine-banded armadillo neonates do not leave their nest before a few weeks after birth (McBee and Baker, 1982) and show limited ranges of movement compared to juveniles and adults. This behavioural change could explain such stark differences between neonates and juveniles. However, ranges of axial movements in younger specimens might also be facilitated by the presence of cartilaginous units that could not be taken into account here. The consideration of such cartilaginous parts might have resulted in higher values of integration and disparity in early stages of development. The timing of this elevated integration actually corresponds with the development of the xenarthrous articulations that ossify postnatally (Hautier et al., 2018) and may facilitate interactions among these tightly interlocking joints (Oliver et al., 2016).

As vertebrae acquire diverging complex morphologies associated with specialized features through development (Johnson and O’Higgins, 1994), disparity was expected to increase during ontogeny. Our results confirm this expectation with whole-column disparity increasing through development and vertebrae occupying more morphospace (Fig. 4). From neonates onward, all three regions display an increase in disparity, concurrent with the differentiation of more specialized structures (metapophysis, anapophysis, diapophysis, and zygapophysis). Following early foetal development, the three vertebral regions occupy different parts of the morphospace and follow divergent paths through development (Fig. 4), which is reflective of their different functions (Oliver et al., 2016). This pattern seems to be already well established in juveniles and to a lesser extent in neonates (Fig. 4).

The drop in disparity that occurs from foetuses to neonates was more unexpected, and is not reflected in the PT analyses. The region-specific disparity analyses enabled us to look more closely at this pattern (Fig. 5B). We observed that the drop in disparity is found almost entirely in the anterior thoracic vertebrae, which suggests that vertebrae in this region display distinctive shapes early during development. Several studies (e.g. Head and Polly, 2015; Jones et al., 2018b) also found that the first thoracic vertebra of *Mus musculus* is morphologically regionalised with the post-atlanto-axial cervical series, which is partly consistent with the results of our regionalisation analyses with two specimens showing the first two anterior thoracic vertebrae gathered in their own region (specimen G, Fig. 3A; specimen D, Fig. 3B). Such a definite grouping between posterior cervicals and anterior thoracics is, however, not predicted for *Dasypus*, as their cervical vertebrae are totally lacking the neural spines that are so prominent in thoracic vertebrae, including the most anterior ones. Alternatively, this developmental pattern of disparity might be attributable primarily to differences in ossification timing, and find explanation in the peculiar ossification sequence of the thoracolumbar vertebrae in *Dasypsus* (Hautier et al., 2011; Fig. 7). While all early foetal vertebrae display a rather simple and homogenous saddle shape, they ossify at different times. In particular, the anteriormost sets of thoracic neural arches ossify before their posterior neighbours (Fig. 7). The early foetal peak in disparity of anterior thoracics could be due to a relatively more advanced development in younger specimens reflective of differential timings of ossification, which might be mitigated later in development when all vertebrae are well ossified. Such an explanation would also account for the diminishing effect of this pattern through ontogeny, when the more posterior thoracic vertebrae start exhibiting more derived morphologies.

### Rates of growth

Our findings indicate that the regionalisation of the vertebral column may stem from different vertebral growth rates. These results have to be taken cautiously as growth rates might be variable through ontogeny. Differences might exist between prenatal and postnatal growth, with vertebrae showing slow prenatal growth but fast postnatal growth (Bergmann et al., 2006), or regions growing more quickly than others but for a shorter period of time (Reichling and German, 2000). Our dataset prevents us from identifying specific bursts of growth through ontogeny or differences in growth rates between stages. Few data exist on the role of growth in the evolution of the axial regionalisation (Bergmann and Russell, 2001; Bergmann et al., 2003, 2004, 2006; Jones et al 2014). Bergmann et al. (2006) exhibited differential growth rates between vertebrae and vertebral regions in rats, with individual vertebrae showing increasing growth craniocaudally. Jones and German (2014) concurred with such a pattern of craniocaudally increasing growth and showed that growth of a particular vertebra is a function of its position within the vertebral column. However, all these previous attempts focused on the development of the centrum, whose shape is more easily traceable using X-ray radiography on longitudinal data. Our results based on neural arches showed rather contrasted results, with the neural arches of the posterior thoracic region growing slowest, and the anterior thoracic and lumbar growing at increasing rates, craniad and caudad respectively. Such differences suggest that distinct developmental pathways contribute to axial skeleton regulation of the neural arches and centra, which was confirmed by previous analyses of modularity in mammals (e.g. Hautier et al., 2010; Randau and Goswami, 2017b). Several studies also identified different genetic influences on neural arch and centrum ossification (Koseki et al., 1993; Wallin et al., 1994a; Peters et al., 1999), with Pax-1 mutant mice being characterized by persistent neural arches in vertebrae where centra are absent (Koseki et al., 1993). In terms of function, slower growth rates in the middle of the dorsal column may reflect increasing relative stiffness in response to sagging forces that occur midway between the limbs (Smit, 2002).

Henry et al. (2007) showed that vertebral regions in naked mole rats respond differently to systematic growth hormones to drive differentiation. We argue that regional differences in the ossification sequence and growth patterns might be determined by regionalisation, which in turn shape morphology. Indeed, some consistent patterns seem to emerge between differential growth rates and ossification timings (Fig. 7). The vertebral region that displays the slowest growth rates corresponds to the last region to ossify its neural arches, which comprises the meeting point between an ossification that spreads caudally from the anterior thoracic region and another that spreads cranially from the lumbar region. In the mouse, the delayed ossification region also matches with the posterior limit of HoxA4 and is near the anterior limit of HoxA9 expression, so that Hautier et al. (2014) suggested that ossification may be another aspect of axial skeleton phenotype influenced by *Hox* regulation. Mutations in *Hox* complexes lead to modification in vertebra identity, as defined by vertebra shape and connectivity to ribs/girdles at the end of foetal development in the mouse (Wellik and Capecchi, 2003). However, while *Hox* genes express very early during development, no *Hox* gene has yet been shown to directly affect vertebral growth rate or timing of ossification, and the direct effects of the *Hox* code on the later processes of vertebral development have yet been elucidated. In mouse and chick models, these later processes include the determination of ventral somite cells into sclerotome cells (Neubuser et al., 1995; Muller et al., 1996), the migration of sclerotome cells, condensation and differentiation of chondroblasts (Wallin et al., 1994b; Peters et al., 1999; Makino et al., 2013), the chondrocyte differentiation and hypertrophy (Crean et al., 1997; Makino et al., 2013), and the endochondral and perichondral ossification through differentiation of osteoblasts from sclerotome cells and calcification (Hermann-Kleiter et al., 2009; Makino et al., 2013). All these events involve different sets of developmental genes. In essence, differential growth timings and rates might constitute an integral part of vertebral identity, and could be influenced by *Hox* genes or by their downstream patterning. However, several developmental factors such as mechanical loadings are also known to influence vertebral growth, and the duration and rate of growth of a particular vertebra might only reflect the pattern of growth of surrounding tissues (O’Higgins et al., 1997).

## CONCLUSION

Here we applied a developmental perspective to the regionalisation of the axial skeleton of *D. novemcinctus* and showed that vertebrae adopt their unique morphology and become consistently regionalised early during development. Although vertebral morphologies transition gradually, the number of vertebral regions seems to be conserved during development despite drastic changes in morphology. The distinction between pre-diaphragmatic non-xenarthrous vertebrae and post-diaphragmatic xenarthrous vertebrae may constitute the more dominant shift, as demonstrated by our regionalisation and trajectory analyses, while the classic division of the trunk between thoracic and lumbar might represent a more secondary shift and relate more to ribs than vertebrae themselves. By tracing the development of vertebral morphologies from foetal to adult stages, we showed that vertebral morphotypes and axial modularity are likely influenced by variations in size, developmental tempo and growth rates. Further comparisons, notably between the centra and the neural arches, are needed to understand how these growth patterns are genetically mediated.

## Acknowledgments

We thank Judy Chupasko, Mark Omura and Mark Renczkowski [Museum of Comparative Zoology, Harvard University (MCZ)] for all of their help during the course of this project, Frank M. Knight (University of the Ozarks) and Benoit de Thoisy (Institut Pasteur, Cayenne, French Guiana) for donating armadillo carcasses, and several colleagues in the MCZ for helping with experimental setup and data collection (Robert Kambic, Brianna McHorse, Blake Dickson and Hanna Barnes). A version of this paper was submitted by J.D.O. in partial fulfilment of the Erasmus Mundus Master Programme in Evolutionary Biology.

## Author contributions

Concepts and approach were developed by J.D.O., L.H. and S.E.P. Experiments and data analysis was performed by J.D.O., L.H., K.E.J. and S.E.P. The manuscript was prepared by J.D.O. and L.H., and edited by K.E.J. and S.E.P. prior to submission.

## Supplementary data

**Figure S1.**
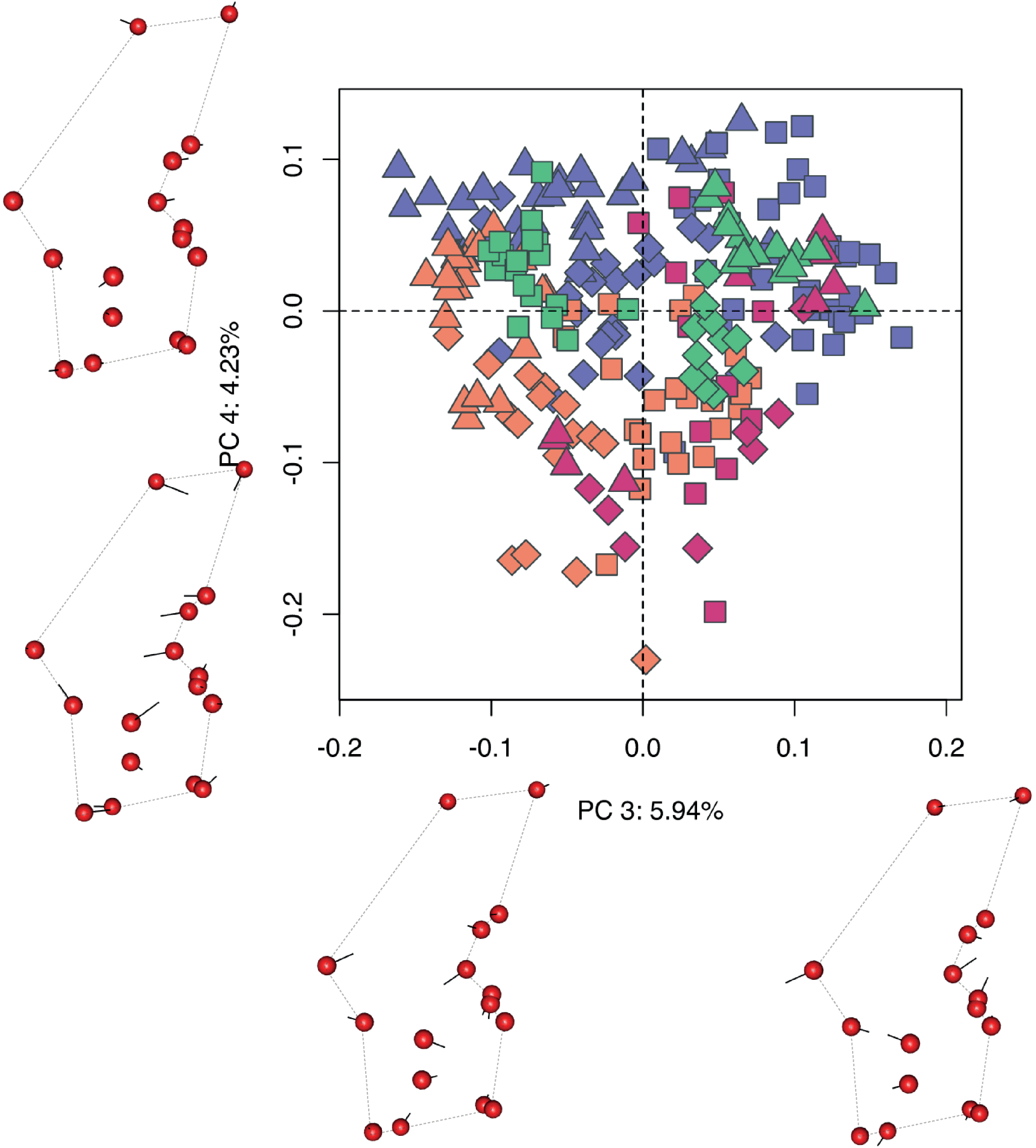
Vertebral morphospace as identified by the third and fourth principal components and associated morphological transformation. Dotted lines represent the outline of neural arches; vectors (black segments) underline the main directional changes. Symbol shapes characterize vertebral regions: squares are anterior thoracics, diamonds are xenarthrous thoracics, and triangles are lumbars. Colors indicate developmental stages: foetuses are in orange, neonates in green, juveniles in pink, and adults in violet.

